# Gating mechanism of human N-type voltage-gated calcium channel

**DOI:** 10.1101/2021.07.08.451561

**Authors:** Yanli Dong, Yiwei Gao, Shuai Xu, Yuhang Wang, Zhuoya Yu, Yue Li, Bin Li, Bei Yang, Xuejun Cai Zhang, Daohua Jiang, Zhuo Huang, Yan Zhao

**Affiliations:** National Laboratory of Biomacromolecules, CAS Center for Excellence in Biomacromolecules, Institute of Biophysics, Chinese Academy of Sciences, Beijing 100101, China; State Key Laboratory of Brain and Cognitive Science, Institute of Biophysics, Chinese Academy of Sciences, 15 Datun Road, Beijing, 100101, China; College of Life Sciences, University of Chinese Academy of Sciences, Beijing 100049, China; State Key Laboratory of Natural and Biomimetic Drugs, Department of Molecular and Cellular Pharmacology, School of Pharmaceutical Sciences, Peking University Health Science Center, Beijing, 100191, China; Laboratory of Soft Matter Physics, Institute of Physics, Chinese Academy of Sciences, Beijing 100190, China; IDG/McGovern Institute for Brain Research, Peking University, Beijing, 100871, China

## Abstract

N-type voltage-gated calcium (Ca_V_) channels mediate Ca^2+^ influx at the presynaptic terminals in response to action potential and play vital roles in synaptogenesis, neurotransmitter releasing, and nociceptive transmission. Here we elucidate a cryo-electron microscopy (cryo-EM) structure of the human Ca_V_2.2 complex at resolution of 2.8 Å. This complex structure reveals how the Ca_V_2.2, β1, and α2δ1 subunits are assembled. In our structure, the second voltage-sensing domain (VSD) is stabilized at a resting-state conformation, which is distinct from the other three VSDs of Ca_V_2.2 as well as activated VSDs observed in previous structures of Ca_V_ channels. The structure also shows that the intracellular gate formed by S6 helices is closed, and a W-helix from the DII-III linker is determined to act as a blocking-ball that causes closed-state inactivation in Ca_V_2.2. Collectively, our structure provides previously unseen structural insights into fundamental gating mechanisms of Ca_V_ channels.

## Introduction

Voltage-gated calcium channels (Ca_V_ channels) are a type of essential mediators to convert action potential into influx of Ca^2+^ ions, a crucial secondary messenger to regulate a variety types of cellular events, such as muscle contraction, secretion of the neurotransmitters, cell division, differentiation, and apoptosis ^1-4^. Ca_V_ channels are generally categorized into two groups according to their activation threshold, i.e., high-voltage activated (HVA) and low-voltage activated (LVA) Ca_V_ channels. Based on their sequence homology, Ca_V_ channels in mammals contain ten members, which are further classified into three subfamilies (Ca_V_1, Ca_V_2, and Ca_V_3) and six types (L-, P/Q-, N-, R-, T-types) ^5-7^. The N-type Ca_V_ channel, also called Ca_V_2.2, is a HVA channel, belongs to the Ca_V_2 subfamily (containing P/Q-, N-, and R-type), and is exclusively expressed in the central and peripheral neurons ^8^. The Ca_V_2.2 is predominantly located in the pre-synaptic terminals and mediates the Ca^2+^ influx that triggers neurotransmitter release at the fast synapses ^9,10^. Dysfunctions of Ca_V_2.2 channels alter neuronal functions and lead to diseases, such as myoclonus-dystonia-like syndrome ^11^. Moreover, the Ca_V_2.2 channel plays a critical role in spinal nociceptive signaling, and thus it has become an important drug target for chronic pain treatment.

Molecular mechanisms of the Ca_V_ channels have been studied extensively for decades, including recent structural studies of L-type Ca_V_1.1 isolated from rabbit skeletal muscle ^12-15^ and human T-type Ca_V_3.1 ^16^. These studies revealed structural features of the Ca_V_ channels at their inactivated state with all the voltage sensing domains (VSD_I_−VSD_IV_) adopting an activated ’up’ conformation. The structures of the Ca_V_1.1 complex also elucidate the assembly of Ca_V_ channel with auxiliary β and αδ8 subunits. A range of ligands were determined in complexes with Ca_V_1.1 or Ca_V_3.1, and these structural studies provide molecular bases for ligand recognition and facilitate further rational drug development targeting Ca_V_ channels. Despite these advances in structural studies on the Ca_V_ channels, more structural insight is desirable to fully understand the molecular mechanism of the Ca_V_2.2 channel from structural untouched Ca_V_2 subfamily, because not only it shares low sequence identity with the available structures from the other two subfamilies, but also some fundamental mechanistic questions remain to be addressed. For example, whereas the available structures of Ca_V_ channels are featured with activated VSDs ^12-14,16^, these VSDs at their resting state appear to be required to fully understand the channel gating mechanism. Moreover, the Ca_V_2.2 harbors a distinct closed-state inactivation during repolarization ^17-19^ and is modulated by phosphatidylinositol 4,5-bisphosphate (PIP_2_) molecules ^20-25^. These underlying molecular mechanisms need further investigation to elucidate.

Here, we purified the recombinant human Ca_V_2.2 (also called α-subunit) in complex with the β1 and α2δ1 subunits, and determined their complex structure using single particle cryo-EM method. This structure reveals the first view of asymmetric activation of the four VSDs with the VSD_II_ at its resting state, presumably locked by a PIP_2_ -like lipid molecule. Furthermore, we identify a unique α-helix that blocks the intracellular gate of Ca_V_2.2, contributing to the close-state inactivation of the Ca_V_2.2. These results enable us to gain significant novel insights into the activation and inactivation mechanisms of human Ca_V_ channels.

## Results and discussion

### Structure determination and architecture of the Ca_V_2.2 complex

To investigate the architecture of the N-type Ca_V_2.2 complex, we co-expressed human N-terminal truncated Ca_V_2.2, full-length wild type (WT) α2δ1, and full-length WT β1 in HEK 293 cells. To monitor the expression of the complex during purification, the Ca_V_2.2 was fused with a C-terminal GFP-Twinstrep tag. In order to enhance the expression level, the intrinsically disordered N-terminal region (residues 1−64) was truncated, and we denote this construct as Ca_V_2.2^EM^. We carried out whole-cell patch clamp experiments in HEK 293T cells to characterize the channel properties of both human WT full-length Ca_V_2.2 and Ca_V_2.2^EM^ constructs, in the presence of auxiliary β1 and α2δ1 subunits. The Ca_V_2.2^EM^ complex shows undistinguishable gating properties to the full-length Ca_V_2.2, in terms of voltage-dependent activation and steady-state inactivation (Fig. 1a-1b; Extended Data Fig. 1a-1b). Thus, the Ca_V_2.2^EM^ construct was subjected to further structural and functional studies. The purified Ca_V_2.2^EM^-α2δ1-β1 complex (Ca_V_2.2 complex) displaying monodisperse peak on both SEC and SDS-PAGE further confirmed that all three subunits were present in the complex sample (Extended Data Fig. 1c-1d).

**Figure 1.**
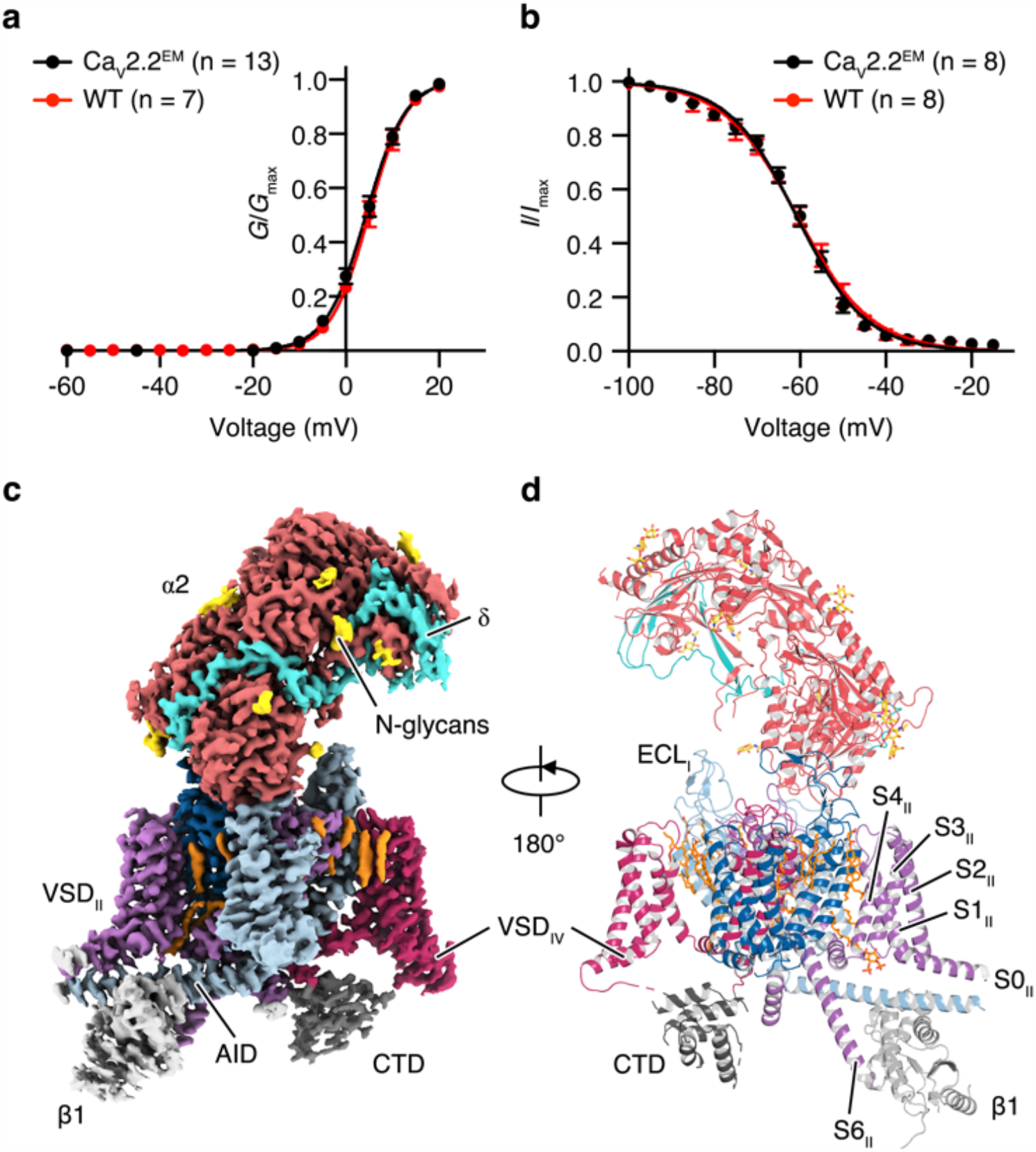
Architecture of the Ca_V_2.2 complex. **a**. Normalized conductance-voltage (*G*/*V*) relationship and the Boltzmann fits for the N-terminal truncated Ca_V_2.2^EM^ complex and full-length wild type (WT) Ca_V_2.2 complex. HEK 293T cells expressing the complex were stimulated with 200-ms depolarizing pulses between −60 mV and 50 mV in steps of 10 mV from a holding potential of −100 mV. **b**. Steady-state inactivation of the WT Ca_V_2.2 complex and the Ca_V_2.2^EM^ complex. Cells were stepped from a holding potential of −100 mV to pre-pulse potentials between −100 mV and -15 mV in 5 mV increments for 10 s. Black, Ca_V_2.2^EM^ complex; red, full-length wild-type Ca_V_2.2 complex. **c**–**d**. The density map and model of the Ca_V_2.2^EM^ complex as seen in parallel to the membrane plane. The α2o1 and β1 subunit, C-terminal domain (CTD), extracellular loops (ECL), alpha-interacting domain (AID), and transmembrane helices S1_II_-S4_II_ in VSD_II_ were labeled. The α1 subunit is colored in light blue (D_I_), violet (D_II_), deep blue (D_III_), magenta (D_IV_), and grey (CTD) respectively. The β1, α2 and o1 subunits are colored in light-grey, tomato-red, and turquoise, respectively. N-glycans is displayed and colored in gold.

To elucidate its architecture, we carried out cryo-EM study of the Ca_V_2.2 complex and obtained the final reconstruction at 2.8 Å resolution for the Ca_V_2.2 complex (Fig. 1c, Extended Data Fig. 2 and Extended Data Table 1). Cryo-EM map of the Ca_V_2.2 is rich of structural features, including densities for side chains, N-glycans, disulfide bonds, and associated lipid molecules, which enabled us to unambiguously de novo build atomic models of the Ca_V_2.2 complex (Fig. 1d and Extended Data Fig. 2e). Of approximately 118 Å × 113 Å × 169 Å in sizes, the structure of the Ca_V_2.2 complex contains α, α2δ1, and β1subunits, and closely resembles the classic shape of Ca_V_1.1.

The α-subunit is composed of four transmembrane domains, D_I_ −D_IV_, which form an ion conducting pore in a domain swapped fashion. Each domain is composed of six helices (S1−S6), of which S1−S4 helices form the voltage sensing domain (VSD). Four VSDs (VSD_I_ −VSD_IV_) encircle the central pore formed by S5 and S6 helices from all four domains. The β1 subunit comprises a Src homology 3 (SH3) domain and a guanylate kinase (GK) domain. The GK domain, through its alpha-interacting domain (AID) helix, interacts with the Ca_V_2.2 (i.e., the α-subunit). Nevertheless, the GK domain was not well resolved in our cryo-EM map, presumably due to conformational heterogeneity (Fig. 1c-1d). The α2δ1 subunit is divided into α2 and δ subunits, which are linked by a disulfide bond (C404-C1059) and associate with the α-subunit by interacting with extracellular loops (ECLs, in particular the loops between S5 and S6 helices) ECL_I_, ECL_II_, ECL_III_, and the S1−S2 loop from VSD_I_. The structure of human α2δ1 in our complex is nearly identical with rabbit α2δ1 resolved in the Ca_V_1.1 complex, with root-mean-square-deviation (r.m.s.d.) of ∼1.1 Å.

### Ion conduction pore of the Ca_V_2.2 complex

The central pore is made up of helices S5 and S6 from all four domains of the α-subunit (Fig. 2a-2b). In each domain, a re-entrant P-loop is located between the S5 and S6 helices and contains two short helices, P1 and P2, connected by a short linker. Four P-loops form a funnel-like shape and line the outer entry to the pore. Four highly conserved glutamate residues, E^314^, E^663^, E^1365^, and E^1655^, form a selectivity-filter ring at the bottom of the funnel, which is one of the narrowest segments of the central pore (Fig. 2c). These four acidic residues, together with surrounding negatively charged residues, create a strong negative electric field strength to attract cations and repel anions. This pore region also functions as a selectivity filter to discriminate Ca^2+^ from other cation ions. We did identify a strong density within the selectivity filter and close to the glutamate cluster (Fig. 2d-2e). Presumably, this piece of density represents a calcium ion, consistent with observations in previous structures of Ca_V_ channels ^12-16^. Thus, on the extracellular side, the S5, S6 helices as well as the P-loops contribute to the formation and stability of the selectivity filter.

**Figure 2.**
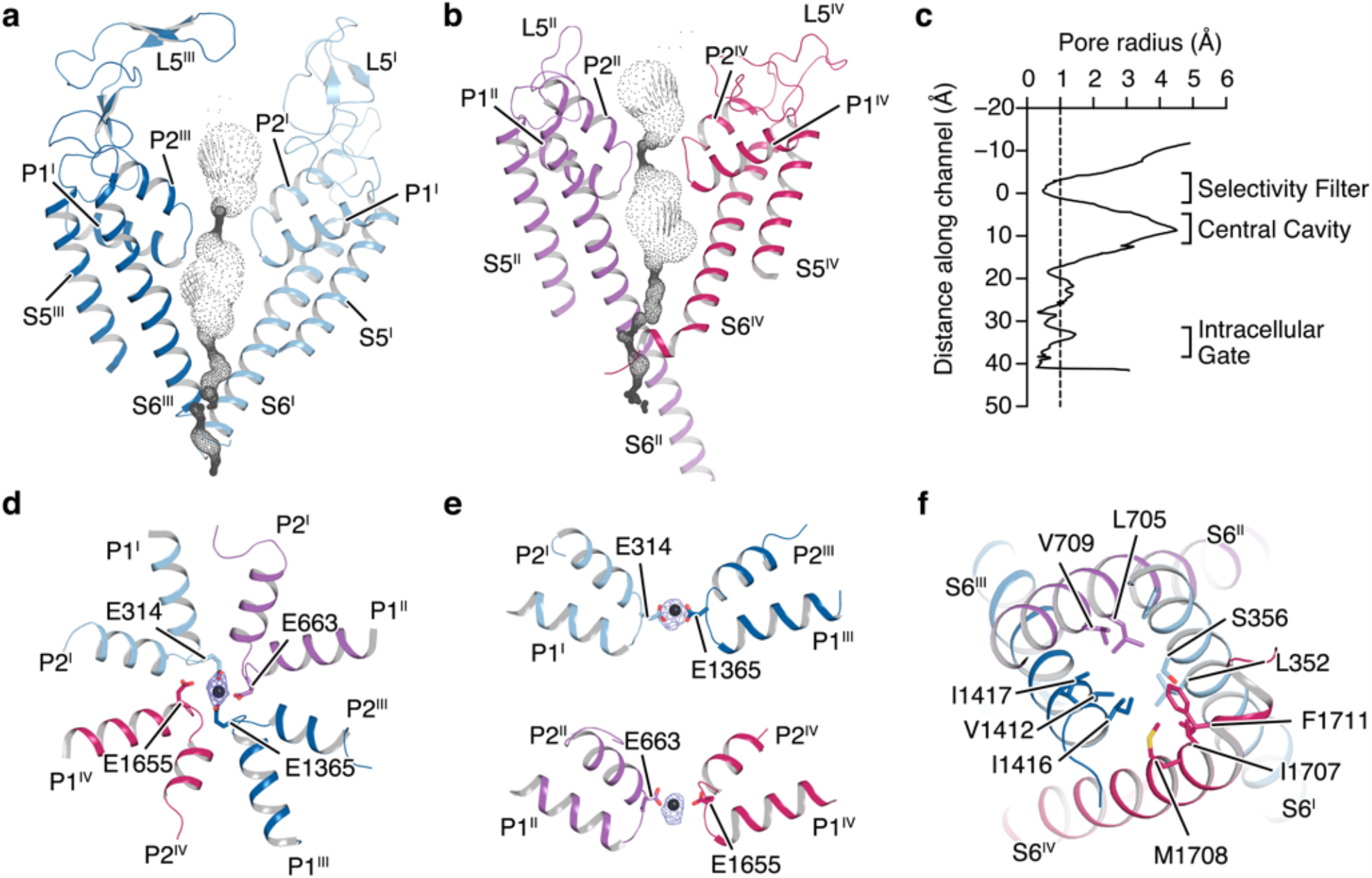
Ion conduction pore of the Ca_V_2.2. **a**–**b**. Ion permeation pathways calculated by the program HOLE were shown in dots. The selectivity filter and S5-S6 helices are shown in cartoon and viewed in parallel to the membrane plane. **c**. Plot of pore radii for Ca_V_2.2 complex. Vertical dashed line marks 1.0 Å pore radius. **d**–**e**. “Top-down” and side-view of the selectivity filter. The selectivity filter ring of four glutamate residues from the four domains of Ca_V_ channel were shown in sticks. A cation ion is shown as a grey sphere, overlaid with corresponding EM density colored in marine. **f**. The intracellular gate formed by four S6 helix viewed from intracellular side. Hydrophobic residues are shown in sticks. The segments from D_I_, D_II_, D_III_ and D_IV_ in the Ca_V_2.2 complex are colored in light blue, violet, deep blue, and magenta, respectively.

On the intracellular side, the pore-lining S6 helices comprise an intracellular gate to control ion permeation (Fig. 2f). The Ca_V_2.2 complex was determined in the absence of membrane potential, and thus is likely to represent a depolarization state. The cytoplasmic ends of the S6 helices are converged and form a hydrophobic seal through a cluster of hydrophobic residues (Fig. 2f). In each of the four domains, the S5 helix is positioned proximal to the S6 helix and connects to S4 of VSD through an amphipathic horizontal helix (termed S4-S5 helix, approx. 16 residues). This S4-S5 helix interacts with the S6 helix, and thus couples S4 movement in responding to depolarization of membrane potential with gate opening.

The central pore of the Ca_V_2.2 channel is further compared with those from the Ca_V_1.1 (PDB ID: 5GJW), and Ca_V_3.1 (PDB ID: 6KZP) channels. The S5-S6 ECLs exhibit nontrivial conformational discrepancy among these structures (Extended Data Fig. 3a, 3b, 3d and 3e). The Ca_V_2.2 α-subunit possesses shorter ECL_I_, ECL_II_, and ECL_III_, but this structural difference does not hamper binding of the α2δ1 subunit. The S1-S2 loops of VSD_II_, also involved in interactions with the α2δ1 subunit, are nearly identical in both Ca_V_2.2 and Ca_V_1.1. Therefore, the α2δ1 subunits in both Ca_V_2.2 and Ca_V_1.1 complexes display similar binding geometry. In contrast, the ECLs of the Ca_V_3.1 channel exhibit remarkable structural discrepancy from the corresponding loops of Ca_V_1.1 and Ca_V_2.2, thus giving rise to incompatibility of the α2δ1 subunit with Ca_V_3.1 (Extended Data Fig. 3b and 3e). Despite of these structural differences occurring at ECLs, the transmembrane helices S5, S6 as well as the selectivity filter of Ca_V_2.2 are superimposable with Ca_V_1.1 and Ca_V_3.1, yielding r.m.s.d of ∼1.8 Å for Ca_V_1.1 (for 332 C_α_ atom-pairs), and ∼1.6 Å for Ca_V_3.1 (for 300 C_α_ atom-pairs), respectively (Extended Data Fig. 3c and 3f). In particular, the intracellular gate formed by the four helix-bundle of S6 helices is nearly identical in all three structures, assuming a closed form (Extended Data Fig. 3g). The S6_II_ helix of the Ca_V_2.2 extends into the cytoplasmic, and is longer than that of the Ca_V_1.1. Consequently, C-terminus of the S6_II_ helix forms a close contact with the β1 subunit in the Ca_V_2.2 complex, which is not observed in the structure of Ca_V_1.1. Furthermore, the cytoplasmic part of the S6 helix bends by 13° compared with that in Ca_V_1.1 (Extended Data Fig. 3h), which may affect gate opening and represent a potential regulatory mechanism of the β subunit.

### Voltage sensor of the N-type Ca_V_ channel

A hallmark feature of the Ca_V_ channel is to open or close the channel in respond to variation of the membrane potential. Surrounding the central pore, the four VSDs play essential roles in converting the electrostatic signal into conformational change of the intracellular gate. Each VSD is composed of S1−S4 transmembrane helices. Each S4 helix contains several positively charged residues, either arginine or lysine, spaced at intervals of three. Ca_V_2.2 is a depolarization-activated channel. Upon depolarization of the membrane potential, the gating charges on the S4 helix move toward the extracellular cavity of the VSD. The two cavities are separated by a highly conserved hydrophobic constriction site. When the gating charges pass through this hydrophobic seal, existing interactions of the gating charges with hydrophilic and negatively charged residues on surrounding helices are disrupted on one side, and new interactions are formed on the other side. The displacement of the S4 helices across the electrostatic field of the membrane potential would induce lateral movement of the S4-S5 linkers, resulting in disengagement between S5 and S6 helices and thus the channel opening.

In our structure, the S4 helices adopt 3_10_ -helix conformation, which is consistent with observations from previous structures of voltage-gated ion channel, including Ca_V_ and Na_V_ channels, allowing the side chains of gating-charge residues to be aligned on the same side of the helix surface. Using the central pore as a reference, superimposition of the structure of Ca_V_2.2 onto that of Ca_V_1.1 shows fairly superimposable VSD_I_, VSD_III_, and VSD_IV_, demonstrating that they are stabilized in the same, presumably activated state (Extended Data Fig. 4a-4e). In contrast, VSD_II_ of Ca_V_2.2 exhibits discernable conformational rearrangement compared with that from the Ca_V_1.1 structure. Taking a closer look at the four VSDs, we found that four, four, and three gating charges are located at the extracellular aqueous cavities of VSD_I_, VSD_III_, and VSD_IV_, respectively, and interact with polar residues residing on nearby S1-S3 helices, consistent with observations from the available structures of Ca_V_ channels ^12-14,16^ (Figs. 3a, 3c, 3d; Extended Data Figs. 4c-4e). Strikingly, in VSD_II_ only one arginine (R578) on the S4 helix is located on the extracellular side of the hydrophobic seal, but the other four conserved gating charges (R581, R584, K587, and K590) are located in the intracellular cavity, indicating that VSD_II_ is stabilized at the resting state. Among the basic residues, K590 is located at a position where S4 becomes unwound, and is completely exposed to the solvent (Fig. 3b). The asymmetric activation of Ca_V_2.2 VSDs suggests that individual VSD may sense membrane potential asynchronously, which is reported to be important for eukaryotic Na_V_ activation and inactivation ^26^.

**Figure 3.**
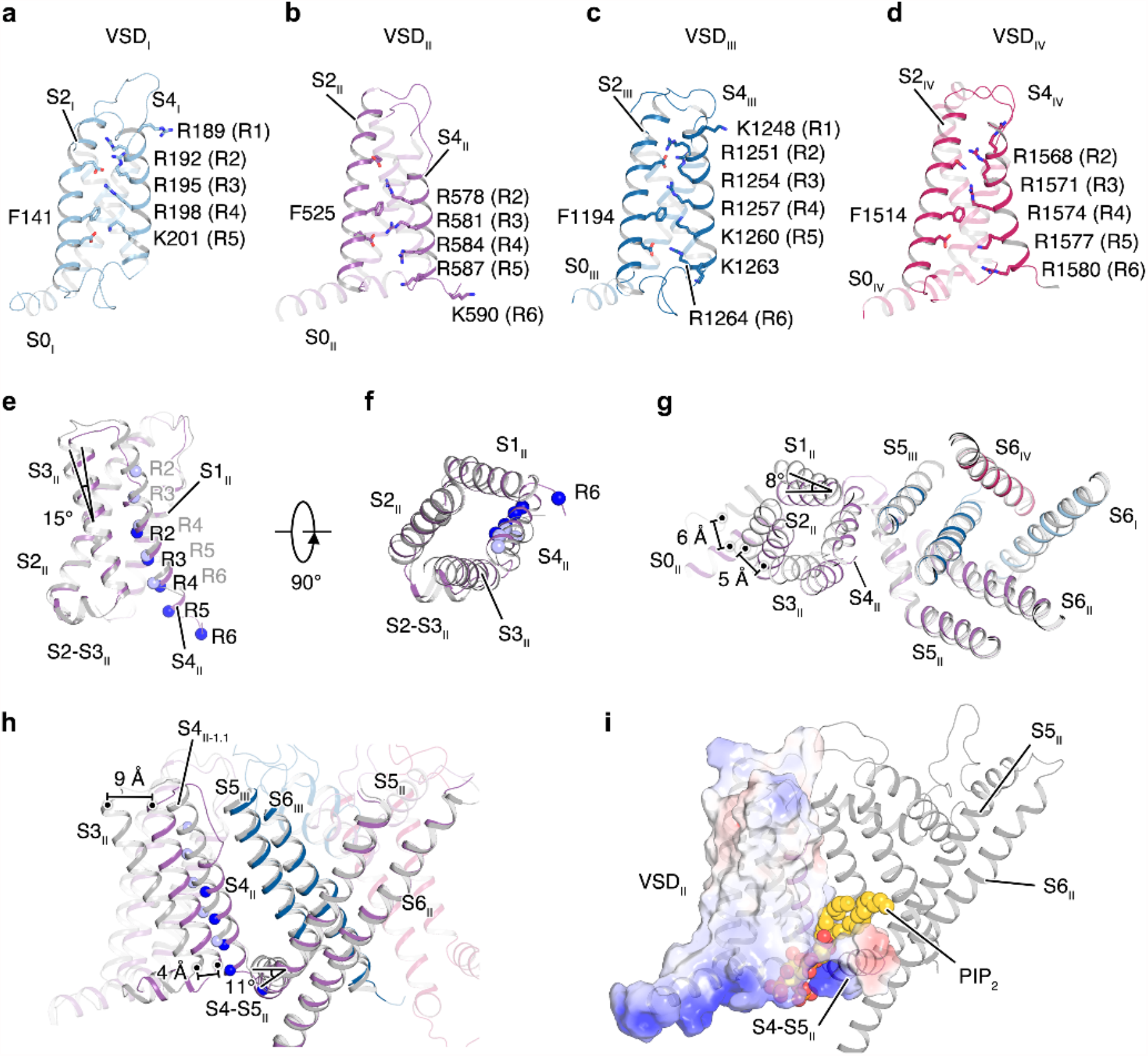
Structural analysis of the voltage-sensing domains. **a**–**d**. Voltage sensing domains from D_I_ (**a**), D_II_ (**b**), D_III_ (**c**) and D_IV_ (**d**) are shown in cartoon. The gating-charge residues (R1-R6) on S4 helix and residues from surrounding helices are shown in sticks. **e**–**f**. Structural comparison of the VSD_II_ at resting state from Ca_V_2.2 complex and activated state from Ca_V_1.1 complex, viewed parallel (**e**) or perpendicular (**f**) to the membrane plane. The gating-charge residues are shown as spheres, colored in deep blue (Ca_V_2.2) or light blue (Ca_V_1.1). **g**–**h**. superimposition of the VSD_II_^R^ and VSD_II_^A^ using S5-S6 helices as a reference, viewed perpendicular (**g**) or parallel (**h**) to the membrane plane. **i**. Domain II of the Ca_V_2.2 complex, overlaid with an electrostatic potential surface on the VSD_II_. The putative PIP_2_ molecular is shown as spheres.

To investigate conformational change of the VSD_II_ of Ca_V_2.2 upon depolarization, we overlaid its structure at the resting state (VSD_II_ ^R^) onto that of Ca_V_1.1 at the activated state (VSD_II_ ^A^), and found that these VSD_II_ structures are highly superimposable in helices S1, S2, and S3 (Fig. 3e-3f). In contrast, the S4 helix slides to the intracellular side in the resting state. Consequently, extracellular halves of S3 helix bends ∼15° toward S4 in the resting state (Fig. 3e). Using the S5 and S6 helices as a superimposition reference, the VSD_II_ undergoes substantial conformational change relative to the central pore between these two states. In particular, in transition from activated state to the resting state, the cytoplasmic sides of both S1 and S2 helices in Ca_V_2.2 rotate toward the central pore (Fig. 3g). Consequently, the intracellular and extracellular terminus of the S3 helices are remarkably displaced approaching to the central pore by 4 Å and 9 Å, respectively. In addition to inwardly sliding, the S4 helices also slightly shift toward the pore domain (Fig. 3h). Interestingly, even though distinguishable conformational change occurs in VSD_II_, the extracellular end of the S1-S2 helix hairpin forms similar interactions with helices S5 and P1 from domain III. It appears that this interaction is stabilized by a cholesterol derivative cholesteryl hemisuccinate (CHS), which is determined in both our structure of Ca_V_2.2 and the structure of Ca_V_3.1 (Extended Data Fig. 4f), consistent with earlier reports suggesting that cholesterol is important to regulating activity of Ca_V_ channels ^27,28^.

Compared with S4 in the activated VSD_II_ from Ca_V_1.1, the S4 helix in the resting state undergoes a sliding movement by two helical-turns, ∼13 Å, towards intracellular side (Fig. 3h). The displacement of S4 is comparable with previous observations in structures of TPC1 channel and Na_V_ Ab channel (Extended Data Fig. 4h-4i) ^29-31^. Except R5 (K590) of Ca_V_2.2, all gating charges are facing to one side, and no rotation about the helix axis is observed, which support a sliding-helix model of voltage-dependent gating ^31,32^. As a consequence of the S4 inwardly sliding, the S4-S5 linker tilted by 11° degree towards intracellular side, and this movement generates more contacts between the S4-S5 linker and S6 helix, and thus stabilize the inner gate at its closed state.

An extra density was identified laying above the S4-S5 linker of domain II in the cryo-EM map, which presumably represents a lipid with two hydrophobic tails and a ‘palm’-like head group. Its head group is located at the intracellular cavity of the VSD_II_ and flanked by S3 and S4 helices (Fig. 3i). Its hydrophobic tails project toward the extracellular side of the membrane (Extended Data Fig. 5a). This lipid appears to act as a plug and would block the S4 helix from upwardly sliding, reminiscent of the PIP_2_ that attenuates opening of the K_V_1.2 channel by interacting with the S4-S5 linker ^33^. We attempted to fit a PIP_2_ into the density. Whereas its hydrophobic tails and inositol ring agree well with the density, the two phosphate groups attached to the inositol ring could not be well resolved (Extended Data Fig. 5a). Considering the PIP_2_ is predominant in the inner-leaflet of the plasma membrane and reported to left shift the activation curve of Ca_V_2.2, which are consistent with our structural observations, we postulate that this lipid molecule is a PIP_2_. However, PIP_2_ molecule was not robustly detected using lipid blot and mass spectrometry methods due to poor yield of the Ca_V_2.2 complex. In a closer look at the PIP_2_ -like lipid binding site, several positively charged residues from helices S0, S4, and S4-S5 linker provide a highly positively charged environment for PIP_2_ binding (Extended Data Fig. 5b-5c). Similar local environment is not observed in the other three VSDs (Extended Data Fig. 5b), implying a mechanism by which PIP_2_ selectively binds with VSD_II_. In addition, the putative inositol ring of PIP_2_ is positioned proximal to the AID helix of the β1 subunit, and this observation is consistent with previous investigations indicating that regulation role of PIP_2_ on the Ca_V_ channel depends on types of β subunits ^34^.

### Inactivation mechanism of the Ca_V_2 channels

The N-type Ca_V_ channel bears a preferential voltage-dependent closed-state inactivation (CSI) ^17,18^, featuring Ca^2+^ insensitive U-shaped inactivation curve, substantial inactivation accumulated during interpulse in a two-pulse protocol and cumulative inactivation during voltage-clamped action potential trains ^17^. In our structure, the intracellular gate is comparable with that in structures of the Ca_V_1.1 and Ca_V_3.1, and appears to be stabilized at a closed state (Extended Data Fig. 3g). Strikingly, an additional α-helix was unexpectedly determined underneath the intracellular gate and flanked by the four S6-helix bundle, forming a 36° angle to the membrane plane (Fig. 4a), and has not been observed in previous structures of Ca_V_ channels. The high resolution cryo-EM map aided us to register the amino-acid sequence to the model. This helix is composed of residues range from A764 to S783, which belongs to the DII-III linker (Fig. 4b). The side chain of W768 points upward into the intracellular gate, forms extensive hydrophobic interactions with residues from S6 helices (Fig. 4b-4c), and thus stabilizes the intracellular gate at its closed state. Thus, we term this 6-turn helix as W-helix. In addition to the tryptophan as a plug inserting into and blocking the intracellular gate, the other residues from the W-helix extensively interact with the S6 helices, including hydrogen-bond and electrostatic interaction, to further strengthen the association of the W-helix with the intracellular gate (Fig. 4b-4c). We speculate that the freshly uncovered W-helix functions as the channel-blocking ball of the “ball-and-chain” model and is important for the stability of the intracellular gate in the closed state, thus contributing to the close-state inactivation of the Ca_V_2.2.

**Figure 4.**
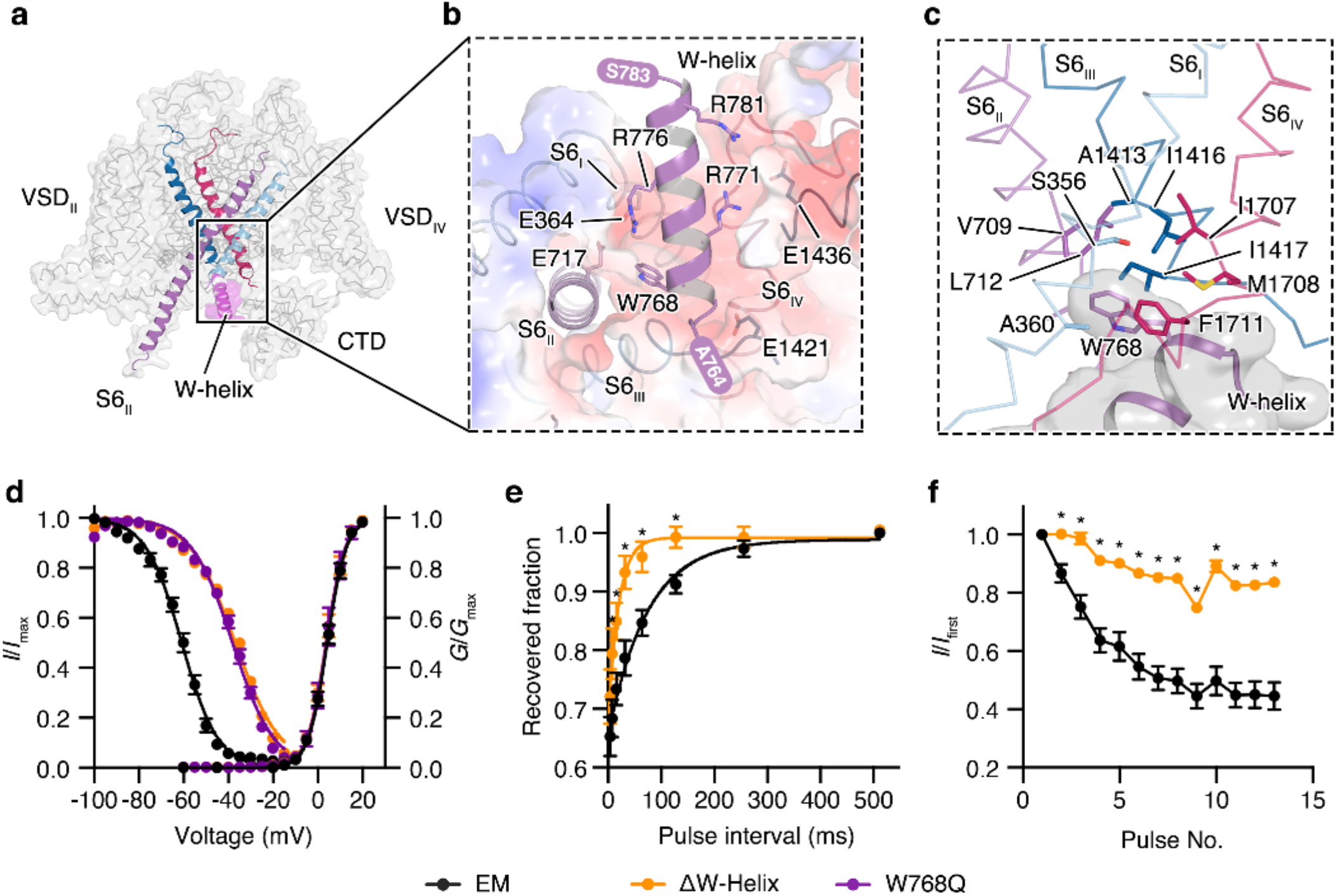
Inactivation mechanism of Ca_V_2.2 complex. **a**. The W-helix is composed of ^764^ARSVWEQRASQLRLQNLRAS^783^ and determined underneath the intracellular gate of the Ca_V_2.2. The Ca_V_2.2 is shown in ribbon and overlaid with transparent surfaces. The S6 helices are shown in cartoon and highlighted. The W-helix is shown in cartoon and overlaid with a transparent surface, colored in purple. **b**. Interactions between the W-helix and the intracellular gate viewed from intracellular side. The critical residues involved in the interactions were shown in sticks. The N- and C-terminus of the W-helix is indicated. **c**. The W768 from W-helix stabilized the intracellular gate at closed state. The S6 helices are shown as ribbon. The W-helix is shown in cartoon and overlaid with a transparent grey surface. **d**. Steady-state activation and inactivation for the Ca_V_2.2^EM^ complex and its mutants. (Activation: EM, n = 13; βW-helix, n = 8; W768Q, n = 6; Inactivation: EM, n = 8; βW-helix, n = 6; W768Q, n = 6) **e**. Recovery of close-state inactivation for the Ca_V_2.2^EM^ complex and its mutants. Cells were held to –40 mV for 200 ms, and then stepped to –100 mV for indicated time delay (4-512 ms), followed by a +10 mV test-pulse (35 ms) (EM, n = 7; βW-helix, n = 8). **f**. Ratio of Cav2.2 channels inactivation calculated from the first spike eliciting maximal current to the other spikes in the AP trains (EM, n = 8; ΔW-helix, n = 6). Black, Ca_V_2.2^EM^ complex; purple, W768Q; orange, ΔW-helix. * indicates *p* < 0.05.

To investigate the functional role of the W-helix and to validate our hypothesis, we designed two mutants, including deletion of the W-helix (Ca_V_2.2^ΔW-Helix^) and a point-mutation variant W768Q (Ca_V_2.2^W768Q^), and performed whole-cell patch clamp recording experiment, with Ba^2+^ as charge carriers. The voltage activation curves of these two mutants were indistinguishable with the WT Ca_V_2.2 complex, suggesting the W-helix might not participate in channel activation. In sharp contrast, the steady-state inactivation curves of these two variants exhibit a depolarizing shift of ∼24 mV (Fig. 4d and Extended Data Fig. 6a-6b), indicating the W-helix and W768 are critical for the CSI. To investigate recovery properties of the CSI, the membrane potential of pre-pulse was held at -40mV, where the channel has not been opened yet. It turns out the recovery of the Ca_V_2.2^ΔW-Helix^ variant is significantly faster than that of WT complex (Fig. 4e and Extended Data Fig. 6c), which further support that the W-helix plays vital roles in CSI of Ca_V_2.2 channel. These results are also in line with a previous functional characterization showing that domain II–III linker is crucial for CSI of Ca_V_2.2 ^19,35^. To further validate our finding in a more physiologically relevant condition, we repetitively activated the Ca_V_2.2 channels with action potential (AP) trains which were recorded in hippocampal CA1 neurons in whole-cell current-clamp configuration. As observed in previous reports ^17,19^, we found that WT Ca_V_2.2 channels exhibited cumulative inactivation in response to AP trains (Fig. 4f and Extended Data Fig. 6d). Interestingly, Ca_V_2.2^ΔW-Helix^ variant inactivated more slowly during the AP train, suggesting that W-helix-regulated close-state inactivation contributed to Ca_V_2.2 channel inactivation during the AP trains, thus may play an important role in short-term synaptic plasticity ^36^. Collectively, W-helix is manifested to be essential for CSI of the Ca_V_2.2 channel. Interestingly, the W-helix is only conserved in the Ca_V_2 subfamily (Extended Data Fig. 7), suggesting that P/Q-type and R-type Ca_V_ channels may adopt the same inactivation mechanism.

## Method

### Whole-cell Voltage-clamp recordings of Ca_V_2.2 channels in HEK 293T Cells

HEK 293T cells were cultured with Dulbecco’s Modified Eagle Medium (DMEM) (Gibco) added 15% (v/v) fetal bovine serum (FBS) (PAN-Biotech) at 37°C with 5% CO2. The cells were grown in the culture dishes (d = 3.5 cm) (Thermo Fisher Scientific) for 24 h and then transiently transfected with 2 μg control or mutant plasmids containing human N-type calcium channel isoform α1B, β1, α2δ1 and GFP using 1.5 μg Lipofectamine 2000 Reagent (Thermo Fisher Scientific). Experiments were performed 12 to 24 hours post transfection at room temperature (21 ∼ 25°C) as described previously ^37^. In brief, cells were placed on a glass chamber containing 105 mM NaCl, 10 mM BaCl_2_, 10 mM HEPES, 10 mM D-Glucose, 30 mM TEA-Cl, 1 mM MgCl_2_, 5 mM CsCl, (pH = 7.3 with NaOH and osmolarity of ∼310 mosmol^-1^). Whole-cell voltage-clamp recordings were made from isolated, GFP-positive cells using 1.5 ∼ 2.5 MΩ fire polished pipettes (Sutter Instrument) when filled with standard internal solution, containing 135 mM K-Gluconate, 10 mM HEPES, 5 mM EGTA, 2 mM MgCl_2_, 5 mM NaCl, 4 mM Mg-ATP, (pH = 7.2 with CsOH and osmolarity of ∼295 mosmol^-1^). Whole-cell currents were recorded using an EPC-10 amplifier (HEKA Electronic) at 20 kHz sample rate and was low pass filtered at 5 kHz. The series resistance was 2 ∼ 4.5 MΩ and was compensated 80 ∼ 90%. The data was acquired by PatchMaster program (HEKA Electronic).

To characterize the activation properties of Ca_V_2.2 channels, cells were held at −100 mV and then a series of 200ms voltage steps from −60 mV to +50 mV in 5 mV increments were applied. The inactivation properties of Ca_V_2.2 channels were assessed with a 10s holding-voltages ranging from −100 mV to −15 mV (5 mV increments) followed by a 135 ms test pulse at +10 mV. To assess the time-dependent recovery from close-state inactivation, cells were depolarized to −40 mV (pre-pulse) for 200 ms to inactivate the Ca_V_2.2 channels, and a recovery hyperpolarization steps to −100 mV were applied for the indicated period (4 ms ∼ 512 ms), followed by a 35 ms test pulse at +10 mV. To analysis the persistent inactivation of Ca_V_2.2 channels following action potential trains, the cells were held at −100 mV and then a physiologically relevant AP train was applied. The AP train used to repetitively activate Ca_V_2.2 channels was obtained from a hippocampal CA1 pyramidal neuron in whole-cell current-clamp mode ^38^. The spike pattern contains 13 action potentials in 2 seconds (mean frequency 6.5 Hz). The percentage inactivation of Ca_V_2.2 channels was calculated from the first spike eliciting maximal current to the other spikes in the AP trains.

All data reported as mean ± SEM. Data analyses were performed using Origin 2019b (Origin Lab Corporation), Excel 2016 (Microsoft), GraphPad Prism 6 (GraphPad Software, Inc.) and Adobe illustrator 2018 (Adobe Systems Incorporated). Steady-state activation curves were generated using a Boltzmann equation.

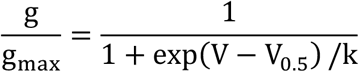

Where g is the conductance, g_max_ is the maximal conductance of Ca_V_2.2 during test pulse, V is the test potential, V_0.5_ is the half-maximal activation potential and k is the slope factor.

Steady-state inactivation curves were generated using a Boltzmann equation.

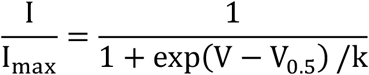

Where I is the current at indicated test pulse, I_max_ is the maximal current of Ca_V_2.2 activation during test-pulse, V is the test potential, V_0.5_ is the half-maximal inactivation potential and k is the slope factor.

Recovery curves from close-state inactivation were the results from seven to nine independent experiments where series of recovery traces from inactivation time points were acquired. The data were fit using a single exponential of the following equation.

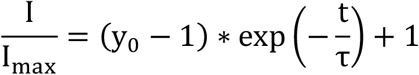

Where I is the current at indicated time delay, I_max_ is the current at 512 ms of time delay, y_0_ is the non-inactivated current at −40mV of pre-pulse, t is the time (the delay between pre-pulse and test-pulse) and τ is the time constant of recovery from close-state inactivation. Statistical significance (P < 0.05) was determined using unpaired Student’s t-tests or one-way ANOVA with Tukey’s post hoc test.

### Expression and purification of human Ca_V_2.2-α2d1-β1 complex

The DNA fragments of human Ca_V_2.2 (UniProtKB accession: Q00975), α2δ1 (UniProtKB accession: P54289) and β1 (UniProtKB accession: Q02641) were amplified from a human cDNA library. The Ca_V_2.2 (full-length wide-type or N-terminal 64-residue truncated), α2δ1 and β1 subunits were subcloned into a modified pEG BacMam vector. A superfolder GFP (sfGFP) and a Twin-Strep tag were tandemly attached at the C-terminus of the Ca_V_2.2 for monitoring protein expression and affinity purification, respectively. The components of Ca_V_2.2 complex were co-expressed in HEK 293F cells using the Bac-to-Bac baculovirus expression system. P1 and P2 viruses of each subunits were obtained using Sf9 insect cells. The P2 viruses were used to infect HEK 293F cells supplemented with 1% (v/v) fetal bovine serum when the cells density reached 2.5 × 10^6^ cells/mL. The infected HEK 293F cells were cultured at 37°C in suspension and 5% CO_2_ in a shaking incubator. 10 mM sodium butyrate was added after 12 hours. The cells were harvested after 48 hours and were stored at −80°C immediately after being frozen in liquid nitrogen.

Cells expressing Ca_V_2.2 complex were resuspended and broken in a purification buffer (20 mM HEPES pH 7.5, 150 mM NaCl, 5 mM β-mercaptoethanol (β-ME), aprotinin (2 μg/mL), leupeptin (1.4 μg/mL), pepstatin A (0.5 μg/mL)) using a Dounce homogenizer. Subsequently, the membranes were collected by centrifugation at 100,000 × g for 1 h and solubilized in the solubilization buffer (20 mM HEPES pH 7.5, 150 mM NaCl, 5 mM β-mercaptoethanol (β-ME), 1% (w/v) n-Dodecyl β-D-maltoside (DDM, Anatrace), 0.2% (w/v) cholesteryl hemisuccinate (CHS, Anatrace)) at 4°C for 2 h with rotation. The insoluble cell debris was removed by centrifugation at 100,000 × g for 1 h. The resulting supernatant was collected and passed through 2 mL Streptactin Beads, which was pre-equilibrated with the wash buffer (20 mM HEPES pH 7.5, 150 mM NaCl, 5 mM β-ME and 0.03% (w/v) glyco-diosgenin (GDN, Anatrace)) and washed with 10 column volumes of wash buffer supplemented with 2 mM ATP and 5 mM MgCl_2_. The Ca_V_2.2 complex was eluted with the wash buffer supplemented with 5 mM desthiobiotin and 1 mM CaCl_2_, and then concentrated in the 100 kDa MW cut-off spin concentrators (Merck Millipore, Germany). For further purification, the concentrated protein sample was subjected to a Superose 6 Increase 10/300 GL gel filtration column (GE Healthcare, USA) pre-equilibrated in the wash buffer supplemented with 1 mM CaCl_2_. The peak fractions between 11.5 mL and 13.5 mL were pooled and concentrated to about 1.8 mg/mL for cryo-EM sample preparation.

### Cryo-EM sample preparation and data collection

A droplet of 2.5 μL of purified Ca_V_2.2 complex was applied on the Quantifoil 1.2-1.3 Au 300 mesh grids glow-discharged for 60 s under H_2_ -O_2_. The grids were then blotted for 4-5 s at 4°C under condition of 100% humidity and vitrified in liquid ethane using a Vitrobot Mark IV.

Cryo-EM data were collected on a 300-kV microscope using a K2 Summit direct electron detector and a GIF Quantum LS energy filter. The slit was set to 20 eV. Movie stacks were acquired at a calibrated magnification of 105,000× in the super-resolution mode, with defocus values ranging from −1.2 to −2.2 μm. The pixel size on motion-corrected micrographs was 1.04 Å. The micrographs were collected under a dose rate of ∼9.6 e^-^/(Å^2^s) and dose-fractioned in 32 frames, yielding a total accumulated dose of ∼60 e^-^/Å^2^.

### Data Processing

A total of 2,300 micrographs were motion-corrected and dose-weighted using MotionCor2 with 5 × 5 patches^39^, followed by CTF estimation using GCTF^40^ and particle picking using both blob picker and template picker in cryoSPARC^41^. Several rounds of 2D and 3D classifications were performed in RELION to remove junk particles ^42^. Initial multi-reference 3D classification generated 6 classes. The class 5 accounts for 26.1% of total particles, displaying discernible structural features, including the secondary structure elements of the α− β1 and α2δ1 subunits. This class were selected and subjected to another round of 3D classification, giving rise to two classes with visible and continuous transmembrane helices. The particles from these two new classes were subjected to 3D refinement, generating a 3.5-Å resolution map. Subsequently, one more round of 3D classification was performed without further particle alignment. Two of the most populated classes were used for further 3D refinement, Bayesian polishing, and CTF refinement in RELION. The particles were then imported into cryoSPARC and subjected to Non-uniform Refinement. The final map was reported at 2.8-Å resolution according to golden standard *Fourier* shell correlation (GSFSC) criterion.

### Model building

The cryo-EM map of Ca_V_2.2 complex showed clear densities for most side chains and N-glycans, which allowed us to reliably build and adjust the model. The homology models of α and α2δ1 subunits were extracted from the structures of *O. cuniculus* Ca_V_1.1 complex (PDB IDs: 5GJW^43^ and 7JPX^44^). The homology models of β subunits were extracted from the structures of *R. norvegicus* Ca_V_ β_2α_ (PDB ID: 1T0J ^45^). All of these homology models were fitted into the cryo-EM map as rigid bodies using the UCSF Chimera ^46^. The resulting model was then manually inspected and adjusted in COOT ^47^, followed by refinement against the cryo-EM map in real space using the phenix.real_space_refine utility. The model stereochemistry was evaluated using the Comprehensive validation (cryo-EM) utility in the PHENIX software package ^48^.

All figures were prepared with ChimeraX or PyMOL (Schrödinger, LLC) ^49,50^.

## Data availability

The three-dimensional cryo-EM density maps of the Ca_V_2.2 complex have been deposited in the Electron Microscopy Data Bank under the accession code EMD-xxxx. The coordinates for the Ca_V_2.2 complex have been deposited in Protein Data Bank under accession code xxxx.

## Author contribution

Y.Z. and D.J. conceived the project. Y.D. carried out molecular cloning and cell biology experiments. Y.D., Y.W., Y.L. and B.Y. expressed, purified protein complex sample and prepared sample for cryo-EM study. Y.D., Y.G. and Y.W. carried out cryo-EM data collection. Y.G. Z.Y.Y. and Y.Z. processed the cryo-EM data and prepared figures. Y.W. and B.L. built and refined the atomic model. Z.H. and Y.Z. designed and S.X. performed electrophysiological experiments. X.C.Z., D.J. and Y.Z. analyzed the structure. Y.Z. prepared initial draft of the manuscript. X.C.Z., D.J., Z.H. and Y.Z. edited the manuscript with input from all authors in the final version.

## Acknowledgments

We thank X. Huang, B. Zhu, X. Li, L. Chen, and other staff members at the Center for Biological Imaging (CBI), Core Facilities for Protein Science at the Institute of Biophysics, Chinese Academy of Science (IBP, CAS) for the support in cryo-EM data collection. We thank Yan Wu for his research assistant service. This work is funded by Chinese Academy of Sciences Strategic Priority Research Program (Grant XDB37030304 to Y.Z. and Grant XDB37030301 to X.C.Z), the National Natural Science Foundation of China (31971134 to X.C.Z., 81371432 to Z.H.) and Institute of Physics, Chinese Academy of Sciences (E0VK101 to D.J.).

## Conflict of interest

All authors declare that there is no conflict of interest that could be perceived as prejudicing the impartiality of the research reported.

## Notes

### Competing Interest Statement

The authors have declared no competing interest.

