## Extended_Data_Figure for "Gating mechanism of human N-type voltage-gated calcium channel"

This file contains Extended Data Figure 1–7 and Extended Data Table 1.

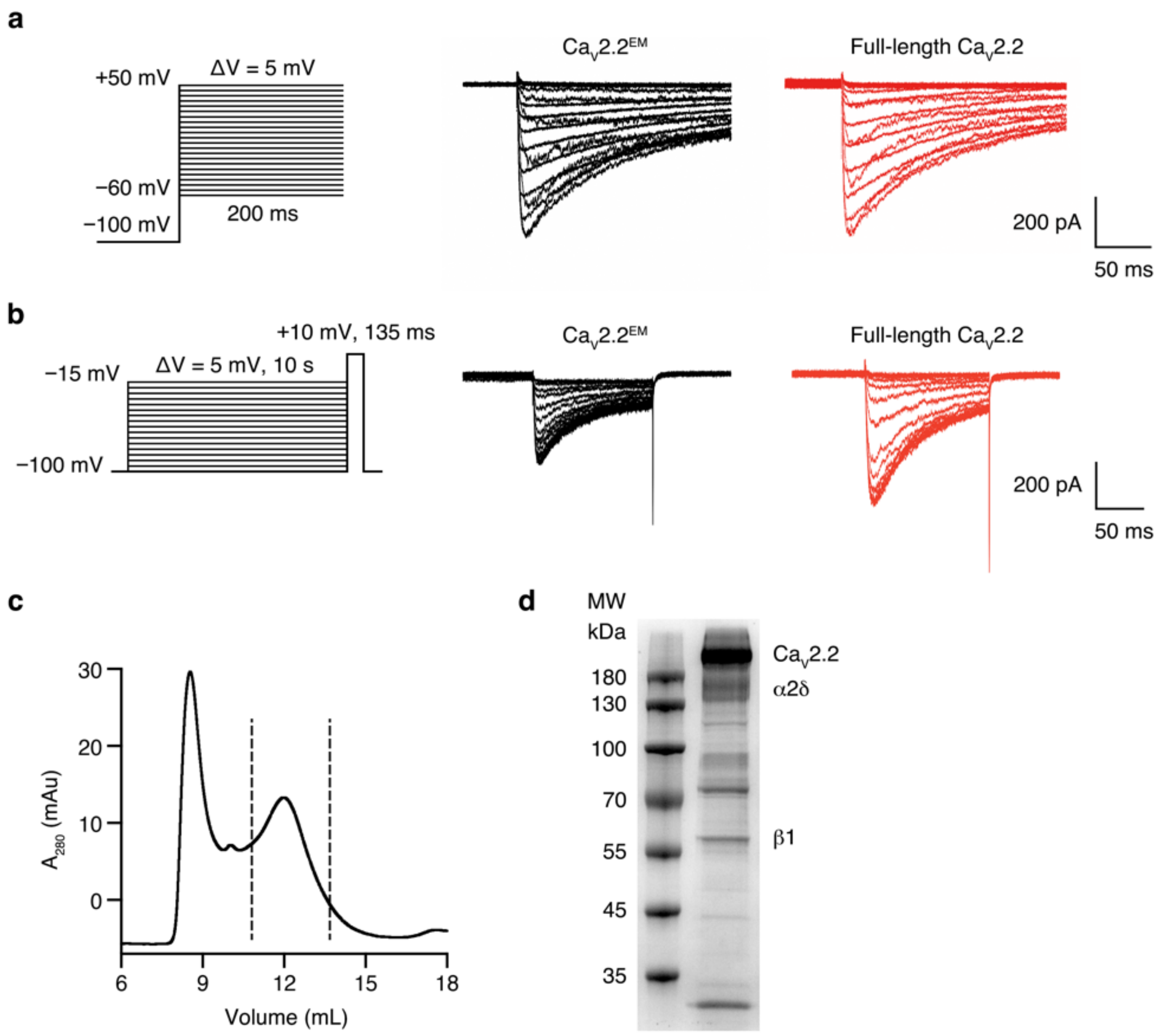

Extended Data Fig. 1 Functional characterization and purification of the Ca$_V$2.2 complex.

**a.** Representative whole-cell voltage-clamp Ca$_V$2.2 current traces obtained from a series of 200 ms voltage steps from –60 mV to +50 mV in 5 mV increments. **b.** Typical whole-cell voltage-clamp Ca$_V$2.2 current traces elicited by a +10 mV test pulse after holding-voltages ranging from –100 mV to –15 mV in 5 mV increments. **c**. Size-exclusion chromatogram (Superose 6 increase) of the purified sample of Ca$_V$2.2 complex. Peak fractions (marked within black dashed lines) were pooled and concentrated for cryo-EM study. **d**. Coomassie-blue-stained SDS-PAGE gel of the purified Ca$_V$2.2 complex. Components of the complex are labeled. The experiments were repeated independently for more than 3 times with similar results.

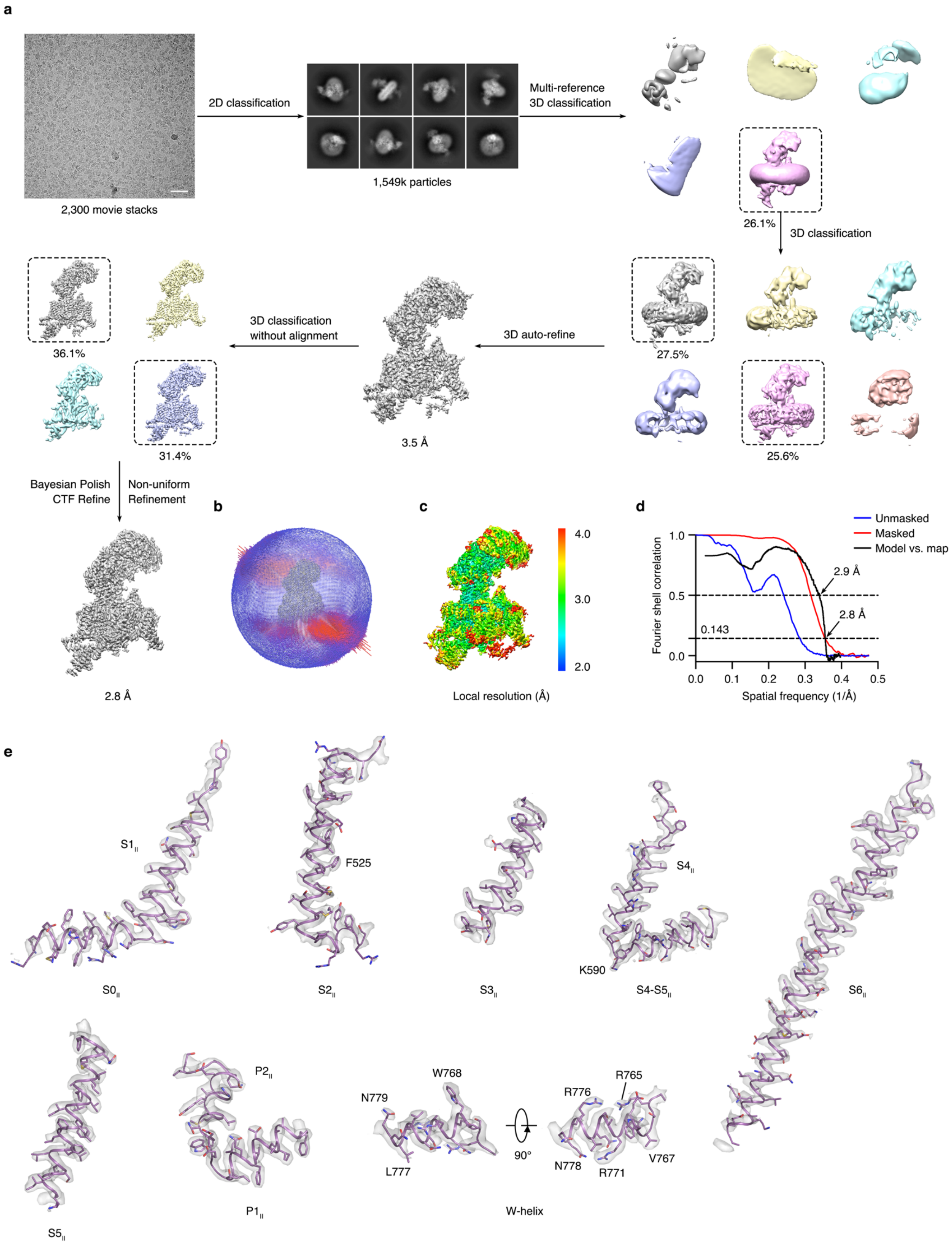
a
2D classification
2,300 movie stacks
1,549k particles
Multi-reference
3D classification
26.1%
3D classification
3D classification
without alignment
3D auto-refine
36.1%
31.4%
27.5%
25.6%
3.5 Å
Bayesian Polish
CTF Refine
Non-uniform
Refinement
2.8 Å
b
c
Local resolution (Å)
4.0
3.0
2.0
d
Unmasked
Masked
Model vs. map
Fourier shell correlation
Spatial frequency (1/Å)
2.9 Å
2.8 Å
0.143
e
S1II
F525
S4II
K590
S0II
S2II
S3II
S4-S5II
S6II
P2II
W768
R776
R765
N779
90°
L777
N778
R771
V767
S5II
P1II
W-helix

Extended Data Fig. 2 Cryo-EM data processing of $Ca_V2.2$ complex.

**a**. Flow chart of cryo-EM data processing. A total of 1,594k particles were picked from 2,300 micrographs. A representative motion-corrected micrograph of this dataset is shown here (Bar = 400 Å). Several rounds of 2D and 3D classifications were conducted to clean particles, followed by Bayesian Polish and CTF Refine to improve image quality. The final map was reported at 2.8 Å according to the GSFSC criterion. **b**. The angular distribution of the final reconstruction. The height of each spike indicated the number of particles in the designated orientation. **c**. Sharpened map of the $Ca_V2.2$ complex, colored according to the local resolution values. **d**. Fourier Shell Correlations (FSC) of the final map of the $Ca_V2.2$ complex, calculated between two independently refined half-maps before (red) and after (blue) post-processing, overlaid with an FSC curve calculated between the cryo-EM density map and the structural model shown in black. **e**. Representative cryo-EM densities of secondary structural elements from domain II and W-helix.

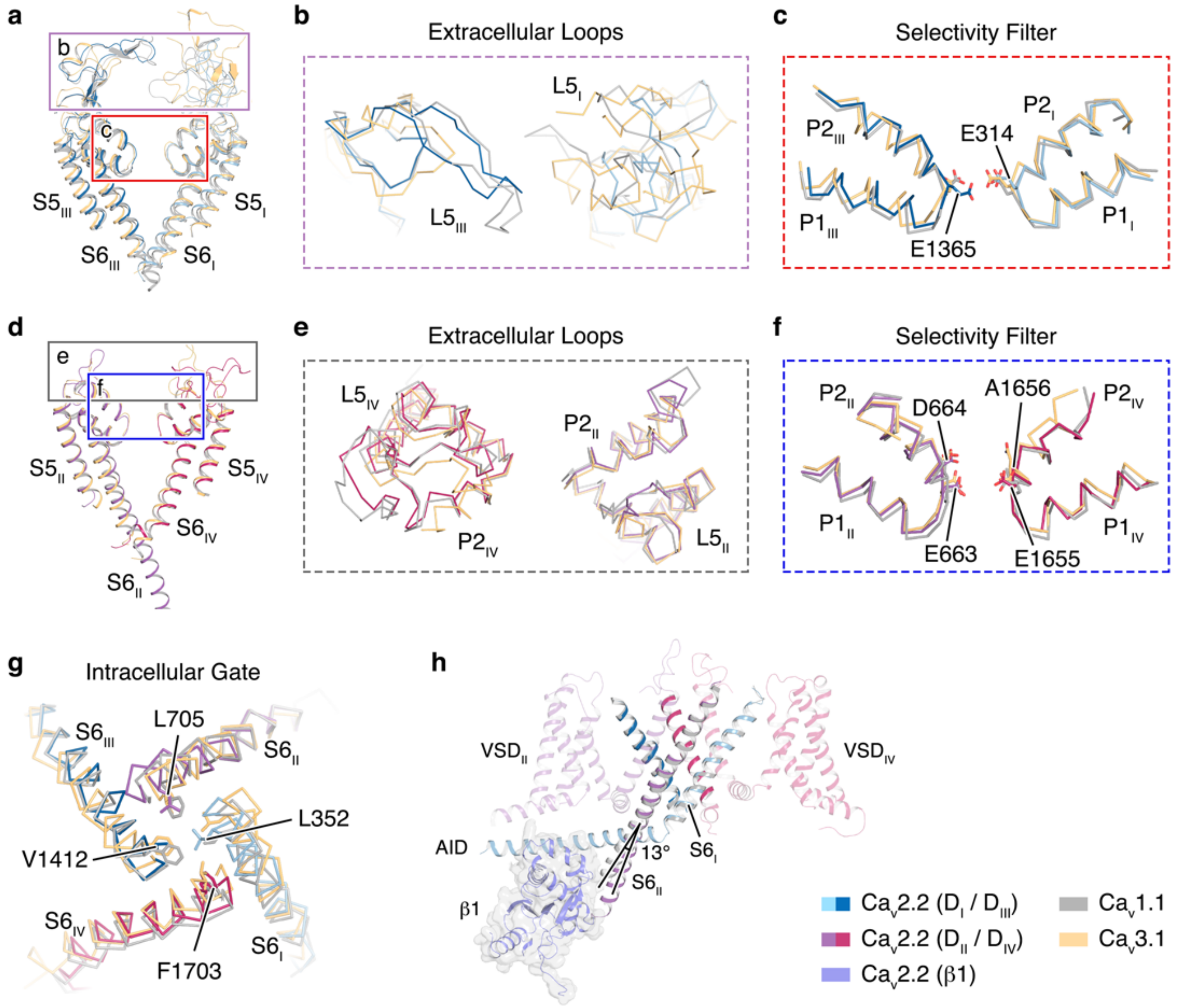

### Extended Data Fig. 3 Structure comparison of the pore domains of Ca$_V$ channels

**a**–**f**. The pore domain segments in the D$_I$/D$_{III}$ (**a**) and D$_{II}$/D$_{IV}$ (**d**), superimposed with Ca$_V$1.1 (grey) and Ca$_V$3.1 (orange). The extracellular loops (**b**, **e**) and selectivity filter (**c**, **f**) are displayed in ribbon. The selectivity-filter ring of four glutamate residues are shown in sticks. **g**. Superimposition of the intracellular gate of the Ca$_V$2.2, Ca$_V$1.1 and Ca$_V$3.1, viewed from the intracellular side. Hydrophobic residues sealing the intracellular gate are shown in sticks. The segments from D$_I$, D$_{II}$, D$_{III}$ and D$_{IV}$ in Ca$_V$2.2 complex are colored in light blue, violet, deep blue, and magenta, respectively. The Ca$_V$1.1 and Ca$_V$3.1 are colored in grey and yellow, respectively. **h**. Structural comparision of the S6$_{II}$ between the Ca$_V$2.2 and the Ca$_V$1.1 (grey), and the interactions among the S6$_{II}$, the β1 subunit, and the AID in the Ca$_V$2.2 complex. The pore domain of Ca$_V$1.1 was superimposed. The β1 subunit in the Ca$_V$2.2 complex was displayed in blue cartoon and overlaid with transparent grey surfaces.

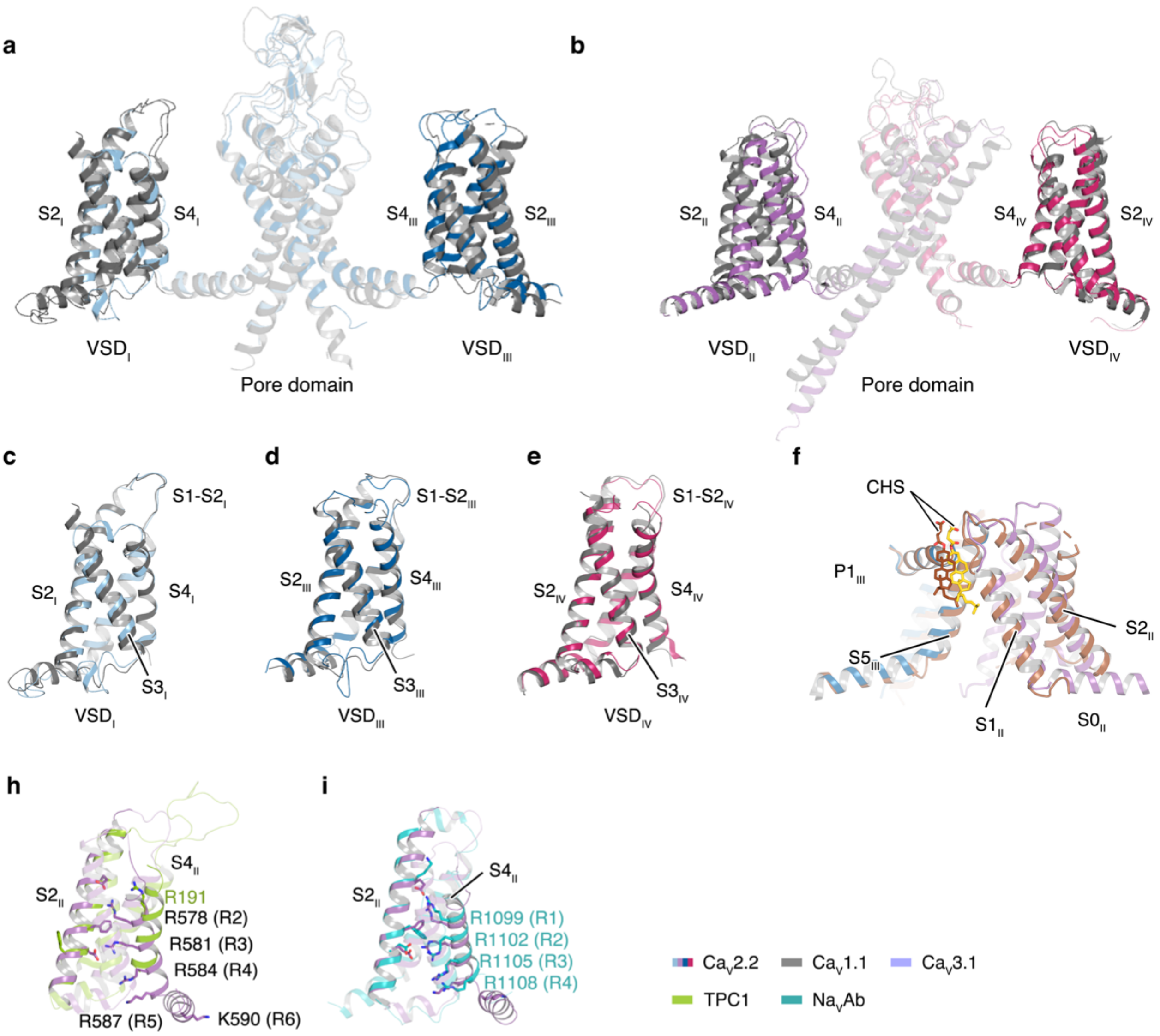

Extended Data Fig. 4 Structural comparison of the voltage-sensing domains

**a**–**b**. The four VSDs of Ca$_{V}$2.2, superimposed with the VSDs of Ca$_{V}$1.1 (grey) using the pore domain as a reference. VSD$_{II}$ in Ca$_{V}$2.2 exhibits a significant structural discrepancy compared to the VSD$_{II}$ in Ca$_{V}$1.1. **c**–**e**. The superimposable VSD$_{I}$, VSD$_{III}$ and VSD$_{IV}$ between the Ca$_{V}$2.2 and the Ca$_{V}$1.1. **f**. The conformational shifts of VSD$_{II}$ relative to the pore domain, superimposed with Ca$_{V}$3.1 (brown). The cholesteryl hemisuccinate (CHS) determined in the Ca$_{V}$2.2 and Ca$_{V}$3.1 were presented as gold and brown sticks, respectively. **h**–**i**. The resting-state VSD$_{II}$ of Ca$_{V}$2.2, superimposed with the resting-state VSDs of TPC1 (green) (**h**) or Na$_{V}$Ab (turquoise) (**i**). The gating-charge residues was shown as sticks. The segments from D$_{I}$, D$_{II}$, D$_{III}$ and D$_{IV}$ in Ca$_{V}$2.2 complex are colored in light blue, violet, deep blue, and magenta, respectively.

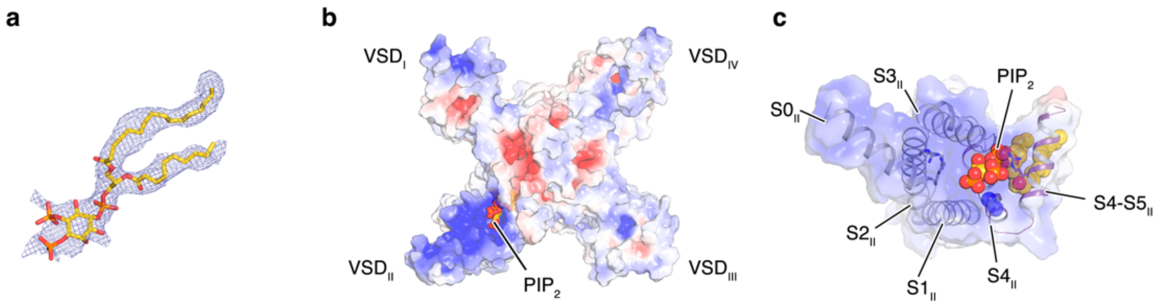

Extended Data Fig. 5 Putative PIP$_2$ molecule associated with the VSD$_{II}$

**a**. The structural model of PIP$_2$, overlaid with the corresponding EM density (blue mesh). **b**–**c**. ‘Bottom-up’ view of the electrostatic surface of the Ca$_V$2.2 $\alpha$1 subunit (**b**) and the VSD$_{II}$ (**c**). The putative PIP$_2$ was displayed as spheres. Positively charged side chains which might contribute to the interactions with PIP$_2$ were shown as sticks.

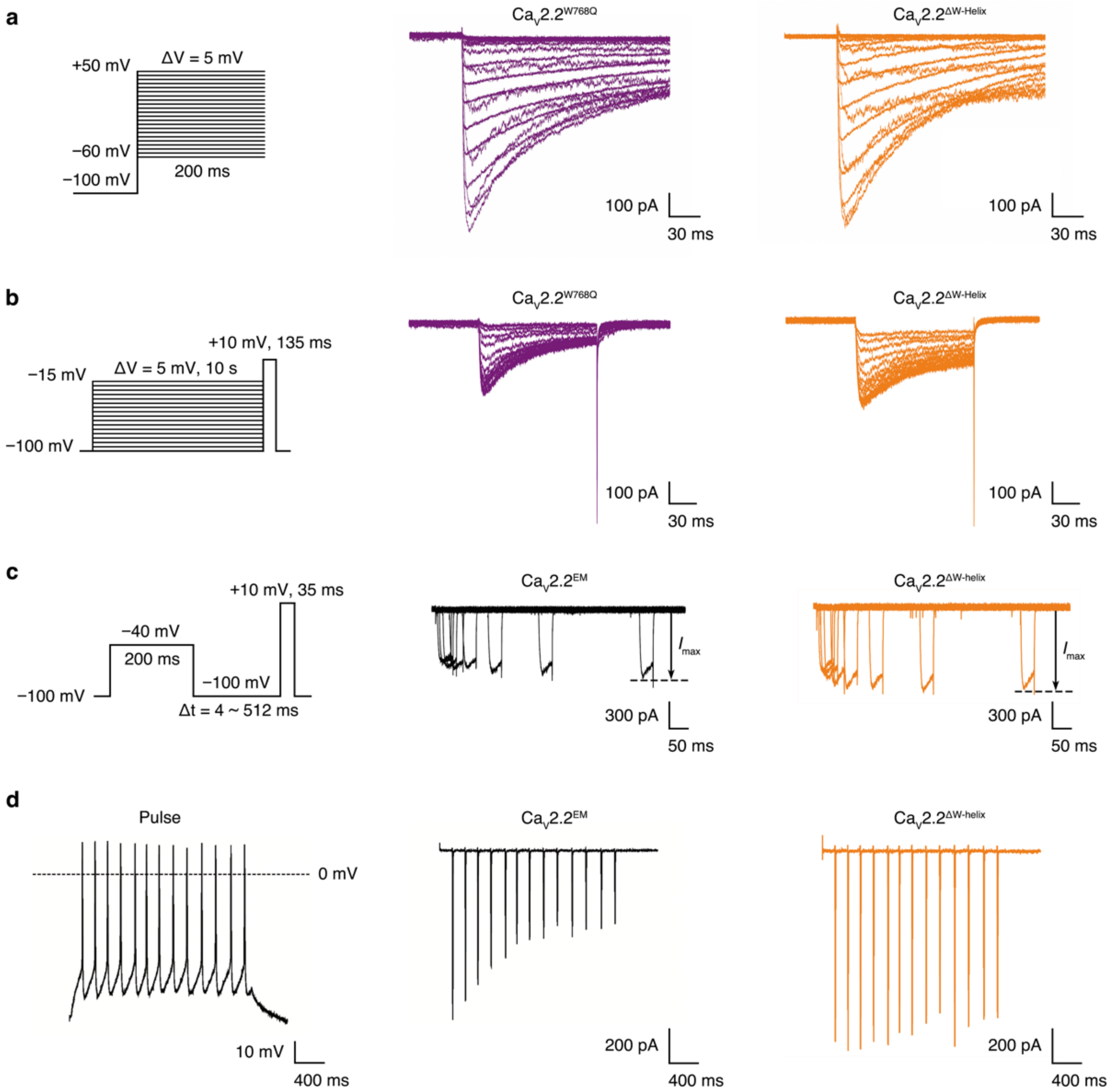

Extended Data Fig. 6 Functional study of the Ca$_V$2.2 complex and mutants

**a.** Representative whole-cell voltage-clamp Ca$_V$2.2 current traces obtained from a series of 200 ms voltage steps from –60 mV to +50 mV in 5 mV increments. **b.** Typical whole-cell voltage-clamp Ca$_V$2.2 current traces elicited by a +10 mV test pulse after holding-voltages ranging from –100 mV to –15 mV in 5 mV increments. **c.** Typical whole-cell voltage-clamp Ca$_V$2.2 currents. Cells were depolarized to –40 mV (pre-pulse) for 200 ms to inactivate the Ca$_V$2.2 channels, and a recovery hyperpolarization steps to –100 mV were applied for the indicated period (4-512 ms), followed by a 35 ms test pulse at +10 mV. **d.** Representative whole-cell voltage-clamp current responses stimulated by action potential trains (left) recorded from a hippocampal CA1 pyramidal neuron in whole-cell current-clamp mode.

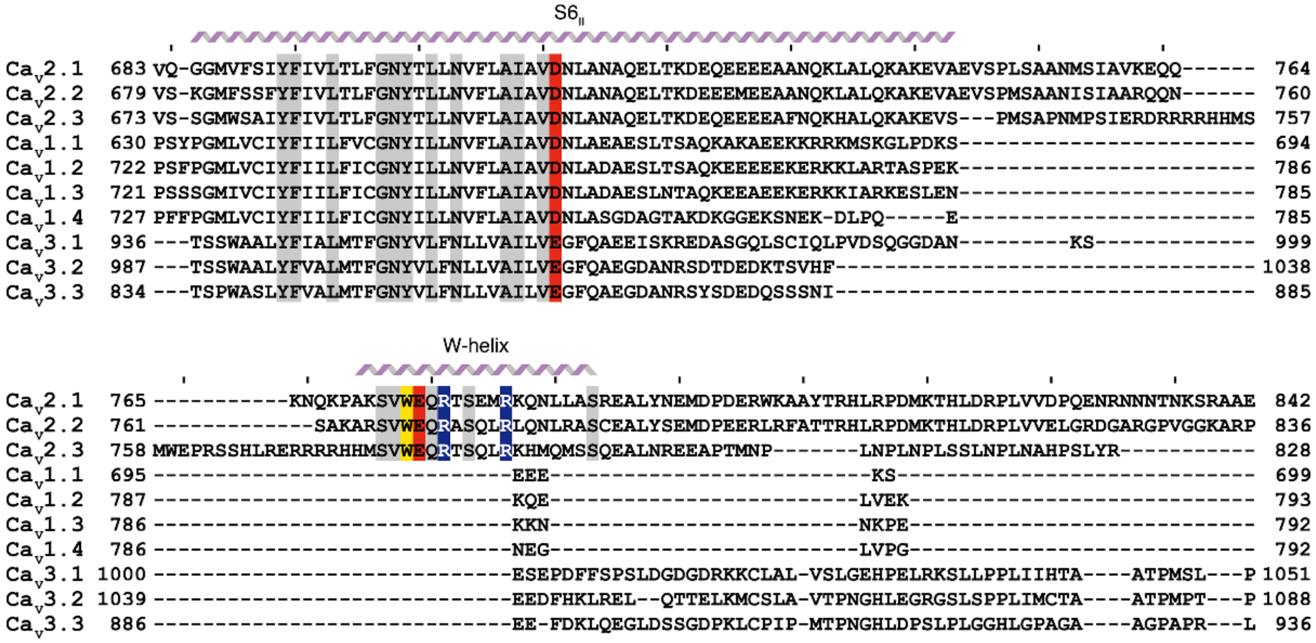

Extended Data Fig. 7 Sequence alignments of the human Ca$_V$ channels.

The sequence alignments of S6$_{II}$ and W-helix among the human Ca$_V$ channels. Positively and negatively charged residues are highlighted in red and blue, respectively. W768 from the W-helix are highlighted in yellow. Other conserved residues are shaded in grey.

**Extended Data Table 1. Cryo-EM data collection, refinement and validation statistics**

| | $Ca_V2.2$ (EMDB-xxxx) (PDB xxxx) |
|---|---|
| **Data collection and processing** | |
| Magnification | 105,000 × |
| Voltage (kV) | 300 |
| Electron exposure ($e^{-}$/Å$^2$) | 60 |
| Defocus range (μm) | −1.2 ~ −2.2 |
| Pixel size (Å) | 1.04 |
| Symmetry imposed | C1 |
| Initial particle images (no.) | 1,584,541 |
| Final particle images (no.) | 253,920 |
| Map resolution (Å) | 2.8 |
| FSC threshold | 0.143 |
| Map resolution range (Å) | 2.0 ~ 4.0 |
| **Refinement** | |
| Initial model used (PDB code) | 5GJW, 7JPX, 1T0J |
| Model resolution (Å) | 2.9 |
| FSC threshold | 0.5 |
| Map sharpening *B* factor (Å$^2$) | −80 |
| Model composition | |
| Non-hydrogen atoms | 19,884 |
| Protein residues | 2,400 |
| Ligands | 41 |
| *B* factors (Å$^2$) | |
| Protein | 58.32 |
| Ligand | 47.84 |
| R.m.s. deviations | |
| Bond lengths (Å) | 0.007 |
| Bond angles (°) | 0.759 |
| Validation | |
| MolProbity score | 2.02 |
| Clashscore | 12.08 |
| Poor rotamers (%) | 0.00 |
| Ramachandran plot | |
| Favored (%) | 93.46 |
| Allowed (%) | 6.54 |
| Disallowed (%) | 0.00 |
